# Inhibition of ice recrystallization with designed twistless helical repeat proteins

**DOI:** 10.1101/2025.03.09.642278

**Authors:** Robbert J. de Haas, Harley Pyles, Timothy F. Huddy, Jannick van Ossenbruggen, Chuanbao Zheng, Daniëlle van den Broek, Ann Carr, Asim K. Bera, Alex Kang, Evans Brackenbrough, Emily Joyce, Banumathi Sankaran, David Baker, Ilja K. Voets, Renko de Vries

## Abstract

Given the repetitive structure of crystalline ice, it’s unsurprising that highly active ice-binding proteins, often with beta-roll structures, also have repeating motifs. Here, we introduce a de novo designed family of ice-binding twistless alpha-helical repeat (iTHR) proteins. Each iTHR protein comprises two planar layers of parallel alpha-helices connected by loops—a structural topology not seen in native ice-binding proteins. The ice-binding helices feature an ordered array of TXXXAXXXAXX motifs, spaced to match the pyramidal {201} and secondary prism plane {110} ice lattice facets, and engineered with a 98.2° turn angle per residue to orient threonines uniformly towards the ice surface. iTHR proteins show high solubility, thermostability, and produce varied ice crystal morphologies depending on their intended target facet. Crucially, iTHRs exhibit ice recrystallization inhibition (IRI) at critical concentrations comparable to those of many native globular ice-binding proteins. Extensive site-specific mutagenesis shows that ice-binding activity in iTHR proteins is robust, remaining largely unaffected by changes in chemical composition. Variation in the repeat number reveals a non-monotonic relationship to IRI activity. X-ray crystal structures of two designs confirm the intended orientation of threonines, uniformly pointing toward the ice surface. The iTHR family provides a versatile platform to systematically investigate the complex structure-activity relationships underlying protein-ice interactions.

## Introduction

Given that the structures and interactions of liquid water and solid ice are so similar, it is remarkable that nature has evolved molecules that bind to ice with high affinity, and often with specificity for particular crystal faces^1,2^. Ice-binding proteins are an important class of ice-binders which have activities ranging from suppressing the growth of small ice-crystals^3^, lowering the freezing point of liquid water^4^, to promoting the nucleation of ice-crystals^5^. Ice crystals possess a highly regular repetitive structure, which is bound by many native ice-binding proteins with geometrically complementary repetitive structures, in particular beta-roll structures^6^. However, many other ice-binding proteins do not have repetitive structures. Additionally, the existence of many ice-binding glycopeptides^7,8^ illustrates that having a precisely folded structure is not a prerequisite for ice-binding.

When designing proteins to bind crystalline surfaces, such as ice, using repeat proteins dramatically reduces the design space to that of a small repeating subunit. This allows large designs to be defined by a small number of degrees of freedom^9–12^. This approach has been used to design helical-repeat proteins with controlled assembly on the mica (001) surface^13^, as well as helical-repeat proteins that promote the nucleation of calcite^14^ or zinc oxide^15^. Recently, it was shown that by a judicious choice of the parameter values, it is possible to specifically design helical-repeat proteins with extended flat surfaces, dubbed <u>T</u>wistless <u>H</u>elical <u>R</u>epeat, or THR proteins^16^. As they contain highly flat repetitive surfaces, similar to those used to bind other crystals, we reasoned that THR proteins should be able to be engineered to bind ice.

Previously, we have shown that the helical TXXXAXXXAXX ice-binding motif, found in ice-binding proteins such as the winter flounder antifreeze protein (wfAFP)^17^, can be incorporated in designed <u>T</u>wist constrained Ice-binding <u>P</u>roteins, or TIPs^18^. These proteins contain an ice-binding alpha-helix featuring TXXXAXXXAXX repeats, two flanking helices which constrain the turn angles of the ice-binding helix, and two loops which connect the helices into a monomeric three-helix bundle^18^. TIPs showed Ice Recrystallization Inhibition (IRI) activity which was maximized when the consecutive motif threonine residues are precisely aligned in the same direction pointing towards the ice surface, by enforcing a helical turn angle of 98.2° in the ice-binding helix (**Fig. 1a**). Similarly to native type-I ice-binding proteins, such as helical wfAFP^17^ and ssAFP^19^, TIPs shape ice into bipyramidal crystals^20^, presumably by binding the pyramidal {201} plane. This behavior of type-I ice binders has been explained by the observation that the threonine residues in the ice-binding (TXXXAXXXAXX)_N_ helices are spaced approximately 16.7Å, a distance that is also present between water oxygens on the pyramidal plane^21–23^. However, this distance also occurs between oxygens on the secondary prism plane {110}, as shown in **Fig. 1b, c**.

**Figure 1.**
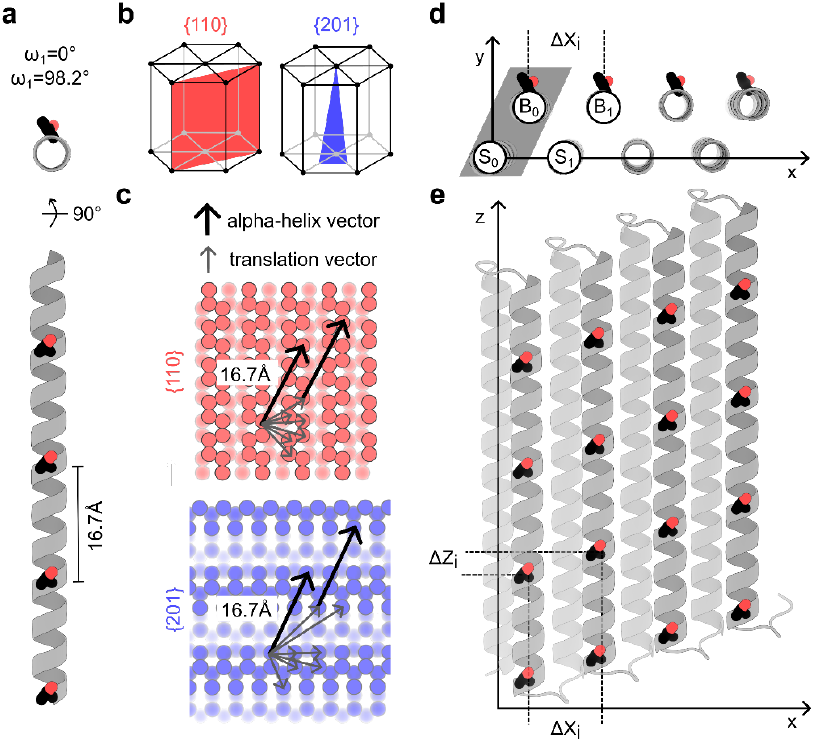
Parametric design of expandable, twist-constrained THRs with threonine matched to oxygen lattice on ice crystal facets. **a)** A 44 residue straight (ω_0_ = 0°) ice-binding helix alpha-helix with 11-mer repeat sequence (TXXXAXXXAXX)_4_ and under-twisted minor helical twist ω_1_ = 98.2°. Threonine residues (sticks) are in perfect alignment, and spacing between each threonine on the 11-mer is spaced 16.7Å. **b)** Examples of ice crystal facet planes (red, blue) on hexagonal ice crystals that have 16.7Å oxygen-oxygen spacing in plane: the secondary prismatic plane {110} and the pyramidal plane {201}. **c)** Crystal slabs of ice facets {110} and {201} with oxygen-oxygen spacing of 16.7Å indicated. Additional vectors indicate potential translations for additional ice-binding helices to multiple rows of oxygen atoms in the plane. **d-e)** Lattice-matched translations in ΔX and ΔZ are applied for *i* number of repeat modules consisting of one or more pairs of stabilising (S_i_) and ice-binding helices (B_i_).

With this in mind, we designed a family of Ice-binding <u>T</u>wistless <u>H</u>elical <u>R</u>epeat (iTHR) proteins, which contain multiple parallel alpha-helices displaying the TXXXAXXXAXX motif spaced to geometrically complement either the {201} or {110} ice surface. A supporting layer of helices constrains the turn angle of the ice-binding helices to 98.2° to align the threonines towards the ice (**Fig. 1a**), and loops connect both sets of alpha-helices into a structure containing tandem repeats. As we will show, the additional design features of the iTHR proteins, as compared to the TIP family of designed ice-binding proteins, allows for designs with more than 1 order of magnitude higher activity than that of the TIP family of ice-binders, having IRI activities that rival that of many native ice-binding proteins.

## Results

### Parametric design of expandable twist constrained iTHRs

We reasoned that a more active family of ice-binding proteins could be obtained if we have arrays of parallel (TXXXAXXXAXX)_N_ ice-binding helices (**Fig. 1d, e**). If we assume the axes of the alpha-helices are along the *z*-axis and the ice-crystal planes to which we want to bind are normal to the *y*-axis, such arrays are characterised by the translations ΔX and ΔZ that relate one ice binding helix to the next one in the array (**Fig. 1c, d**). We intend to design ice-binding THR proteins consisting of two layers of parallel alpha-helices. The top layer contains a first set of binding helices (“B”) that binds to ice and a second set of supporting helices (“S”) that constrain the ice-binding helices to have a turn angle of ω_1_ = 98.2° per amino acid (**Fig. 1d**). Hence, further parameters characterising the iTHR designs are those that relate the position and orientation of the binding helices (B_i_) to those of the supporting helices (S_i_).

To design iTHR ice-binders we started by compiling a set of translations ΔX and ΔZ that allow for threonines on multiple parallel ice-binding helices to interact with repeated sites on the {110} or {201} crystal facets of hexagonal ice (**Fig. 1c**). A set of possible translations ΔX and ΔZ for the ice-binding helices that geometrically match the {110} and {201} facets is given in **Table S1**. We restricted ourselves to values for ΔX and ΔZ within realistic ranges for the design of THR proteins. For example, ΔX values that are too large or too small will not allow for good packing of neighbouring helices.

We decided to design with ice-binding helices of either 22 or 44 residues, allowing for *N* = 2 or *N* = 4 repeats of the putative ice-binding TXXXAXXXAXX repeats. Ice-binding helices B_i_ with *I* = 0, 1, 2, 3 (for the case of *N* = 4), generated by ΔX and ΔZ translations, are constrained by corresponding supporting S_i_ helices that allow for fixing a helical turn of ω_1_ = 98.2° per amino acid in the ice-binding helices B_i_ (**Fig 1d, e**).

For a first set of parametric backbone designs, the repeat unit of the iTHR proteins consists of a single ice-binding helix “B”, and its corresponding supporting helix “S” (grey parallelogram; **Fig. 1e**). For a second set of designs, the repeat unit contains two ice-binding helices and two corresponding supporting helices. This increases diversity in input backbones for improved sampling. Loops of 3-4 residues connecting helices within and between repeats are designed using RosettaRemodel^24^. Because each repeat is symmetrically equivalent, repeats can be propagated to a total of 4 repeats that have internal repeat symmetry.

Next, ProteinMPNN^25^ is used to design amino-acid sequences to populate the looped parametric backbones, while maintaining internal symmetry by tying the identity of residues at the same position within each repeat. The putative ice-binding residues are constrained to follow the TXXXAXXXAXX motif during sequence design, while all other residue identities are chosen by the ProteinMPNN. Residues exposed to the solvent at the capping helices are designed to enhance solubility. Finally, AlphaFold2^26^ and RoseTTAFold^27^ are used to select sequences that are predicted to fold into the designed structures. Designs were selected for experimental screening if they had a mean AlphaFold2 pLDDT score greater than 85, a Cα root-mean-square deviation (Cα-RMSD) of less than 2.5 Å between the AlphaFold2 prediction and the parametric backbone, and a Cα-RMSD of less than 5 Å between the AlphaFold2 and RoseTTAFold predictions.

#### Biochemical characterization of iTHRs

For a first experimental screening of the selected design, a total of 36 designs were cloned with Golden Gate assembly from e-blocks, and proteins were expressed in 4 mL *E. coli* cultures and purified via immobilized metal affinity chromatography (IMAC). 32/36 designs yielded protein bands by SDS-PAGE after purification. Out of these, two {110} binding designs (a-iTHR-110, b-iTHR-110) and two {201} binding designs (a-iTHR-201, b-iTHR-201) were selected for further characterization because they appeared to have high yield, high purity, and the expected molecular weight in SDS-PAGE analysis. Importantly these had large (*N* = 4) ice-binding surfaces. Translations and other design parameters for these four designs are provided in **Table 1**.

**Table 1.** Translations of twist-constrained de novo helical repeat proteins designs. The internal symmetric repeat consists of 1 or 2 ice-binding helices. Translations shown are relative to the first ice-binding helix (B_0_).

| Design | helices<br>per<br>repeat | $\Delta X$<br>($B_0 \rightarrow B_1$) | $\Delta Z$<br>($B_0 \rightarrow B_1$) | $\Delta X$<br>($B_0 \rightarrow B_2$) | $\Delta Z$<br>($B_0 \rightarrow B_2$) | Target facet |
| --- | --- | --- | --- | --- | --- | --- |
| **a-iTHR-110** | 4 | 13.831 | 9.256 | 27.662 | 1.902 | {110} |
| **b-iTHR-110** | 2 | -10.375 | 2.828 | -20.751 | 5.656 | {110} |
| **a-iTHR-201** | 2 | 12.396 | -5.516 | 24.792 | -11.032 | {201} |
| **b-iTHR-201** | 2 | 12.396 | -5.516 | 24.792 | -11.032 | {201} |

For the four selected iTHR designs, AlphaFold2 models were nearly identical to the initial parametrically designed structures, including the location and orientation of the ice-binding threonine residues (**Fig. S1**). All designs contained four threonines per ice-binding helix, except for b-iTHR-110, which had three threonines. The missing threonine was redesigned during loop remodelling to facilitate packing of the loop.

All iTHRs were purified in larger 50 mL cultures and found to be monomeric in Size Exclusion Chromatography (SEC), eluting at the expected retention volumes (**Fig 2b**). Furthermore, all iTHRs showed the expected high alpha-helical content by circular dichroism (CD) and were thermostable (**Fig 2c, d**). Only the a-iTHR-110 design shows gradual unfolding at higher temperatures, but completely refolds after cooling (**Fig. S2**). Mass spectrometry confirmed that all four iTHR proteins had the expected molar mass (**Table S2**).

**Figure 2.**
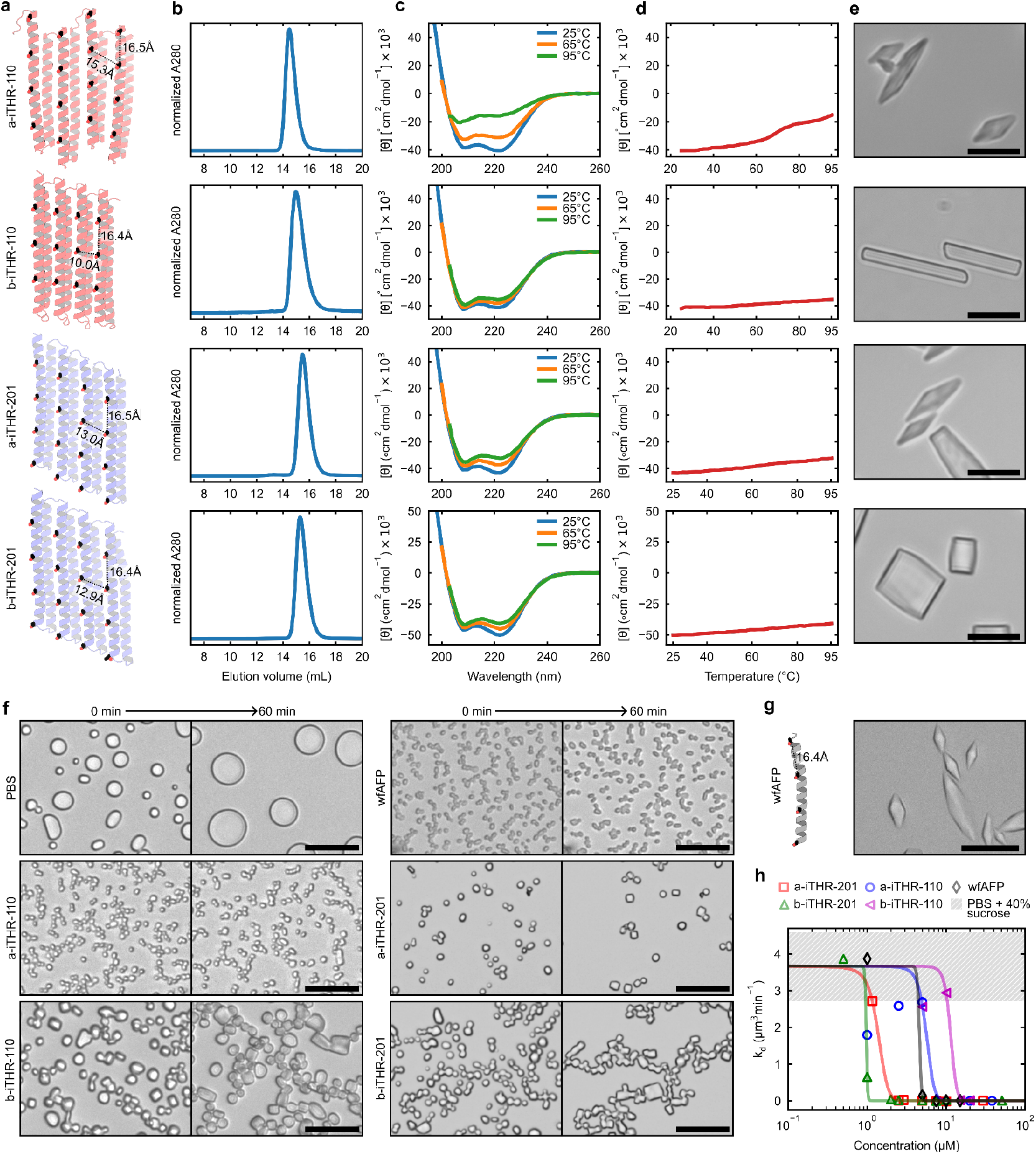
Biochemical characterization and ice-binding activity of iTHRs. **a)** AlphaFold2 models for iTHR designs. Ice-binding threonine residues shown as sticks. The average threonine Cβ-Cβ distances are annotated. **b)** Size-exclusion chromatography of iTHR proteins on Superdex 200 Increase 10/300 column in PBS. **c)** Circular Dichroism spectra in PBS buffer, showing molar residue ellipticity [θ] over wavelength. **d)** Heat ramp at 222 nm, showing molar residue ellipticity [θ] over temperature. **e)** Ice crystal shaping at 113.3 µM a- iTHR-110, 26.0 µM b-iTHR-110, 58.8 µM, a-iTHR-201, and 101.7 µM b-iTHR-201. **f)** Ice recrystallization inhibition screening. Stills at 0 min. and 60 min are shown after incubation at -8°C in PBS + 40% sucrose. Protein concentrations are 38 µM (a-iTHR-110), 26 µM (b-iTHR- 110), 29.4 µM (a-iTHR-201), 50.9 µM (b-iTHR-201), and 15.0 µM (wfAFP). Scale bars: 50 um. **g)** wfAFP (PDB ID: 1WFA) structure (*left*) and ice crystal shaping at 20.0 µM wfAFP (*right*). Scale bar: 25 µm. **h)** Quantitative IRI: ice crystal growth rate (k_d_) vs. protein concentration for a-iTHR-110, b-iTHR-110, a-iTHR-201, b-iTHR-201, and wfAFP. Solid lines are sigmoidal fits to the data. The inflection concentration (C_i_) are 5.6 µM (a-iTHR-110), 11.4 µM (b-iTHR-110), 1.4 µM (a-iTHR-201), 1.0 µM (b-iTHR-201) and 4.5 µM (wfAFP). The mean growth rate in PBS buffer alone was 3.7 ± 0.9 µm^3^/min (n=3 measurements), with the height of the box representing one standard deviation above and below the mean.

#### Characterization of ice-binding activity

Native antifreeze proteins bind to ice facet planes, causing the ice crystals to form distinct shapes. For an initial screening of ice-binding activity, we performed an ice-shaping experiment at high protein concentrations (113 µM, 26 µM, 58.8 µM, and 101.7 µM). An ice-shaping experiment involves nucleating ice crystals in PBS + 20% sucrose containing ice-binding proteins, then slowly cooling the solution further and using microscopy to monitor how the proteins interact with different ice facets to influence the shape and growth of the crystals. We observed a range of distinct crystal shapes, indicating ice-binding activity: a-iTHR-110 formed bipyramidal shapes, b-iTHR-110 produced rectangular and hexagonal crystals, a-iTHR-201 exhibited bipyramidal shapes, and b-iTHR-201 generated blunt-ended bipyramidal crystals (**Fig. 2e**). Full views of the shaping experiment are shown in **Fig. S3**. Compared to the native wfAFP (**Fig. 2g**), a-iTHR-201 showed most similar bipyramidal shapes (**Fig. 2e**).

Ice-crystal shaping is thought to reflect preferences of binding to specific ice-crystal planes^28^. However, the relation is not straightforward since ice-crystal shapes are sensitive to many experimental parameters. Nevertheless, if proteins bind to and stabilise the {110} face one would expect to see rectangular or hexagonal shapes, whereas blunt-ended or bipyramidal crystal shapes as observed for wfAFP are expected for {201} binders^2^. While b-iTHR-110, a-iTHR-201, and b-iTHR-201 align with the observed trends, a-iTHR-110 does not follow this trend. This deviation suggests that the a-iTHR-110 design does not preferentially bind its target facet. Several factors which could contribute to this phenomenon will be explored in the discussion section.

Next, for an initial screening of IRI activity, dispersions of small ice-crystals with and without a fixed concentration ranging from 26 to 113 μM of the iTHR ice-binders in PBS + 40% sucrose were kept at −8°C while monitoring the growth of the ice-crystals. Optical micrographs of the ice-crystal dispersions at 0 and 60 minutes show that ice crystals remain smaller in all iTHR samples than in the control sample without iTHR or the native wfAFP (**Fig. 2f**). To quantify and compare IRI activity, we determined the rate of ice-crystal growth for various concentrations of each of the iTHR designs and wfAFP (**Fig. 2h**).

A key parameter for assessing IRI efficacy is the inhibitor concentration at which the transition occurs from fast recrystallization to slow (if even detectable) recrystallization, marking the curve’s inflection point^3^. We find an inhibitor concentration of Ci = 5.6 μM for the a-iTHR-110 design, while Ci = 11.4 μM for b-iTHR-110 is approximately two-fold higher, indicating inferior performance. Notably, b-iTHR-110 contains one fewer threonine residue per repeat compared to a-iTHR-110 (**Fig. 2a**). In comparison, the iTHR-201 designs demonstrated enhanced mid-point concentrations: a-iTHR-201 with Ci = 1.4 μM and b-iTHR-201 with Ci = 0.97 μM. Both of these designs significantly outperform the wild-type antifreeze protein (wfAFP), which has a Ci = 4.6 μM, and is reported to bind the {201} facet^17^.

#### Importance of residue identity and geometric complementarity on activity

An outstanding question in the literature on ice-binding proteins is to which extent ice-binding is determined by geometric complementarity to the ice-crystal surfaces and to which extent by chemical recognition^29,30^. Our designed iTHR proteins are a unique testing ground to address this issue. We designed steric knockout mutants of a-iTHR-110 in which either the threonine in position 1 of the 11-mer ice-binding motif (TXXXAXXXAXX) (a-iTHR-110-KO.Thr), or the alanine at position 5 (a-iTHR-110-KO.Ala) was mutated to a glutamic acid (**Fig. 3a**). In native antifreeze proteins, often a single or a few steric mutations on the ice-binding surface can significantly disrupt ice-binding activity^31^ and result in a complete loss of TH activity^32,33^. Here, 12 Ala to Glu or 16 Thr to Glu mutations on the ice-binding site reduce IRI activity approximately 3-fold, but surprisingly, significant activity remains for both knockouts: the Ci was 14.6 uM for a-iTHR-110-KO.Ala, and 20.2 uM for a-iTHR-110-KO.Thr (**Fig. 3b**). This suggests that geometric complementarity alone can drive ice-binding, to some extent.

**Figure 3.**
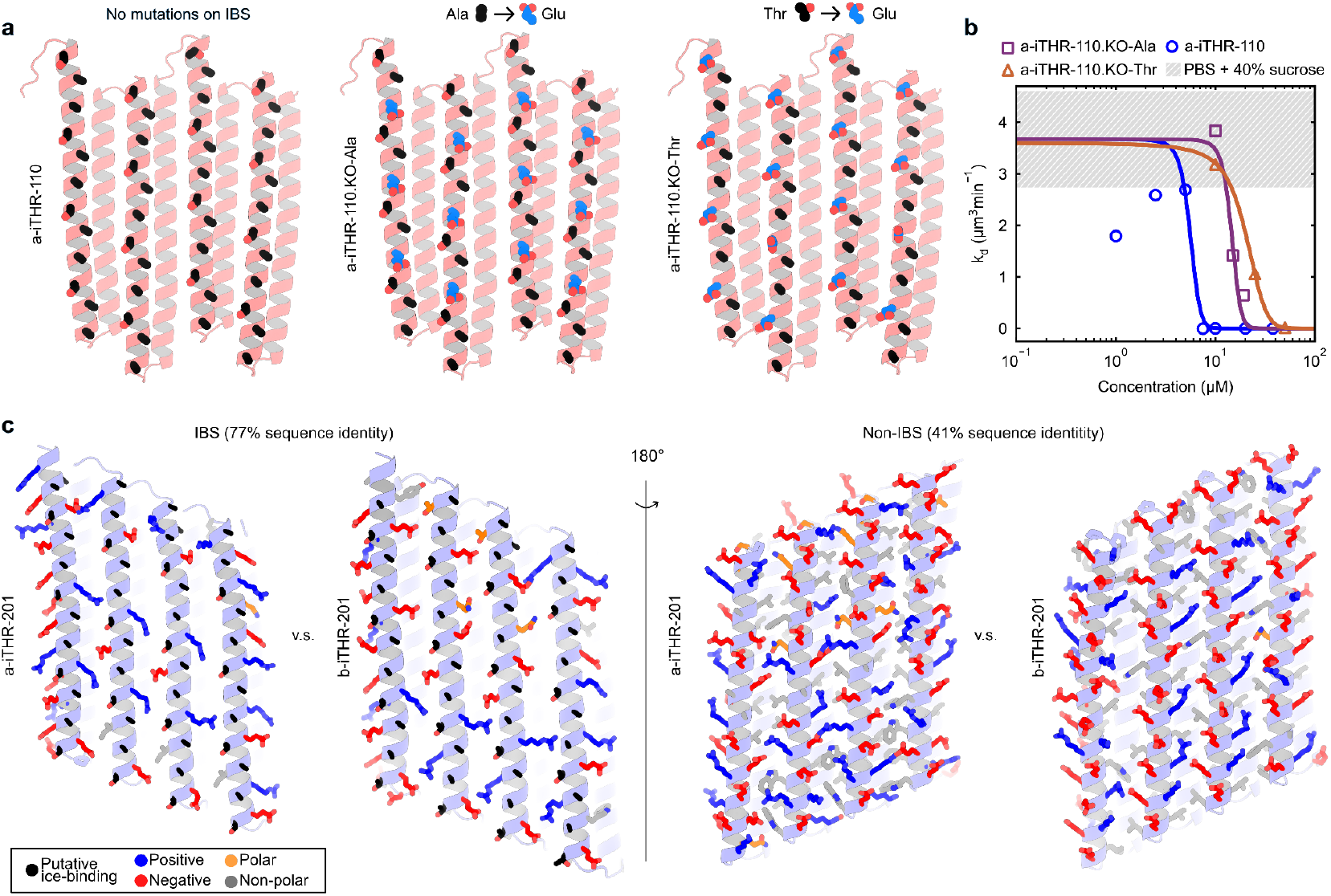
iTHR ice-binding and non-ice binding surface chemistry. **a)** AlphaFold2 models of a-iTHR-110 and knockout mutants a-iTHR-110.KO-Ala and a-iTHR-110.KO-Thr with Ala to Glu or Thr to Glu mutations, respectively. **b)** Quantitative IRI: ice crystal growth rate (k_d_) vs. protein concentration for a-iTHR-110, a-iTHR-110.KO-Ala and a-iTHR-110.KO-Thr. The inflection concentrations are 5.6 µM (a-iTHR-110), 14.6 µM (a-iTHR-110.KO-Ala) and 20.2 µM (a-iTHR-110.KO-Thr). **c)** AlphaFold2 models of the putative ice-binding surface (IBS) and the non-IBS for a-iTHR-201 and b-iTHR-201. Both {201} binders have identical rigid body translations and similar C_i_ (1.4 and 1.0 µM). The surface exposed residues on the ice binding surface (IBS) shows a high degree of sequence similarity of 77%, while sequence identity of surface residues on the non-IBS side was only 41%. Colors of side chains indicate negative (Glu and Asp in red), positive (Lys and Arg in blue), polar (Asn, Gln, Thr, Ser in orange) and hydrophobics (Ala, Val, Ile, Leu, Phe, Trp in grey) residues. The putative ice-binding residues shown in black.

A comparison of a-iTHR-201 and b-iTHR-201 allows for assessing the importance of the surface residues on the non-ice binding side of the proteins: although the two structures exhibit identical backbone displacements in the ΔX and ΔZ directions (**Table 1**), they feature different loops and distinct protein sequences. The alignments of their AlphaFold2 models are very close, with a Ca-RMSD = 1.9 Å (over 1116 atoms). The ice-binding helices on the ice-binding site (IBS) share a high degree of similarity, with a sequence identity of 77% at the ice-binding surface. In contrast, the surface residues on the back sides of IBS show considerable variation, with only about 41% sequence identity (**Fig. 3c**). Given that the C_i_ for both proteins is relatively similar (1.4 µM for a-iTHR-201 and 1.0 µM for b-iTHR-201), this suggests that the identity of the back-side residues is likely not crucial for IRI activity.

### Dependence of activity on iTHR repeats

A key feature of repeat proteins that bind at solid-liquid interfaces is that one can modulate binding affinity by simply changing the number of repeats. This was observed for designed DHRs that adsorb to mica and CaCO_3_ surfaces^13,14^.

To test the impact of the number of repeats on the activity of the iTHRs, we prepared versions of the a-iTHR-201 and a-iTHR-110 proteins with, respectively, twice as many helical repeats (a-iTHR-110×2 and a-iTHR-201×2) and half as many helical repeats (a-iTHR-110×0.5 and a-iTHR-201×0.5) (**Fig. 4a**). IRI activity curves for different iTHR repeat lengths are shown in **Fig. 4b, c**. For the a-iTHR-201 series, the IRI-activity C_i_ of proteins with increasing repeat numbers are 1.4 µM, 5.6 µM, and 1.9 µM. For the a-iTHR-201 series, the C_i_ of proteins with increasing repeat numbers are 0.92 µM, 0.97 µM, and 2.2 µM. For both series we find that increasing the repeat number or IBS did not markedly improve the IRI activity.^34^

**Figure 4.**
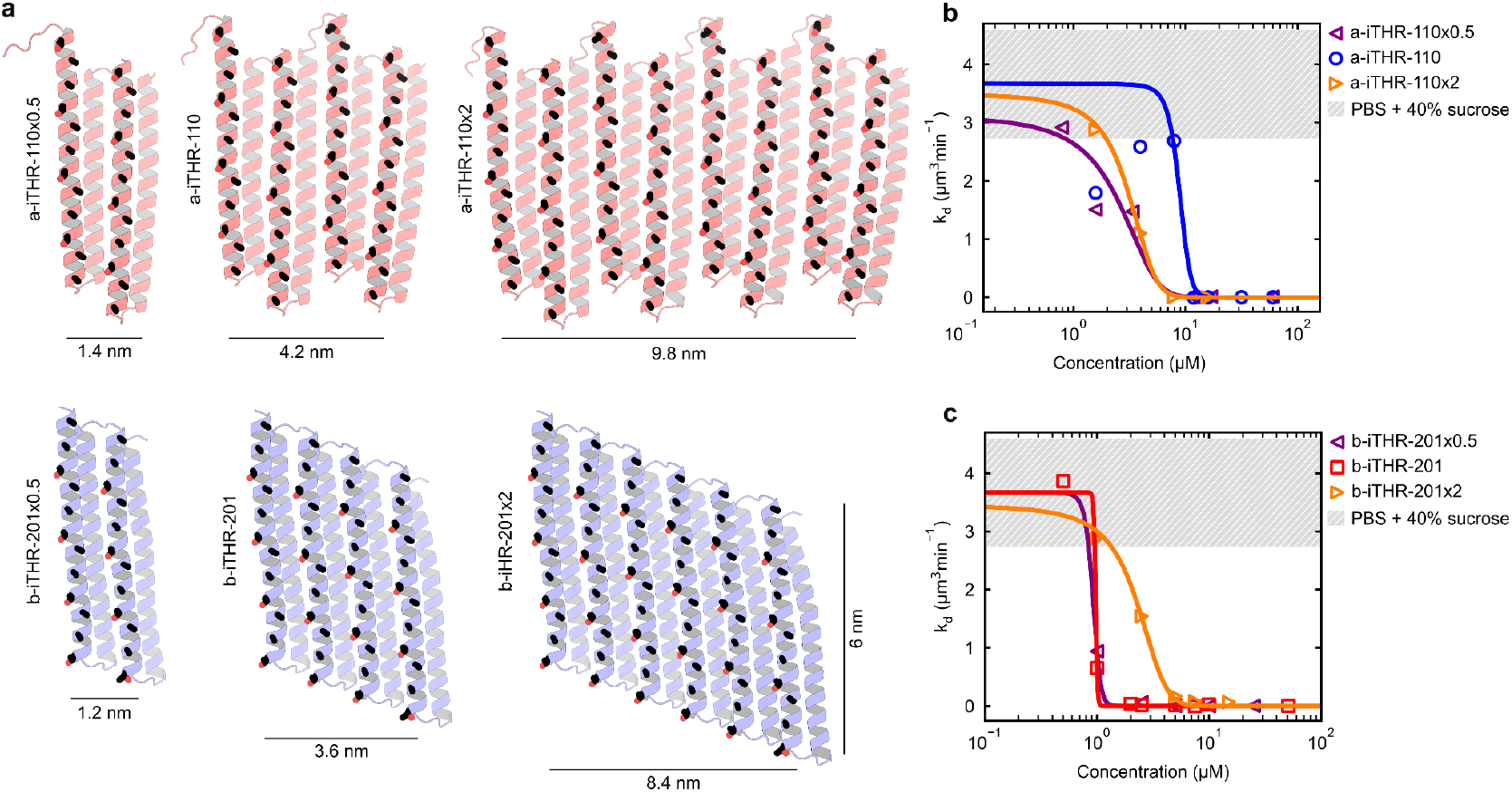
iTHR repeat extension series. **a)** AlphaFold2 models with 0.5x and 2x repeats of a-iTHR-110 and a-iTHR-201. Width of the protein is shown in the figure. Putative ice-binding residues shown in black. The approximate ice-binding surface is 8 nm^2^, 25 nm^2^ and 59 nm^2^ for a-iTHR-110 series, and 7 nm^2^, 22 nm^2^ and 50 nm^2^ for a-iTHR-201 series. **b)** Quantitative IRI: ice crystal growth rate (k_d_) vs. protein concentration for the a-iTHR-110 and a-iTHR-201 extension series. The inhibitory concentrations (C_i_) are 1.4 µM, 5.6 µM and 1.9 µM for a-iTHR- 101×0.5, a-iTHR-110, a-iTHR-110×2, respectively. **c)** Quantitative IRI for the a-iTHR-201 series the C_i_ are 0.92 µM, 0.97 µM and 2.2 µM for a-iTHR-201×0.5, a-iTHR-201, a-iTHR-201×2, respectively.

### Crystallographic structure determination

Finally, to validate experimentally that the iTHR proteins indeed fold precisely into the designed target structures, we attempted to crystalize and solve crystal structures for all four iTHR designs. High resolution structures were obtained for b-iTHR-110 and a-iTHR-201. The a-iTHR-201 structure matches very well both with parametrically designed backbone structure (RMSD = 1.1 Å, **Fig. 5c, d**). The b-iTHR-110 crystal structure is slightly twisted, but otherwise resembles the parametric design backbone (RMSD = 2.5 Å, **Fig. 5a, b**). In both structures, the orientations of the ice-binding threonine and alanine residue are preserved and the ice-binding helix retains the designed alpha-helical turn angle of ω_1_= 98.2° per amino acid of the ice-binding helix. This suggests that our computational design method is able to yield, with high accuracy, the target geometric spacings and residue orientations required for ice-binding. The rotamers of most residues at the core of the protein closely match the design model (**Fig. 5b, d**), demonstrating high accuracy at the sidechain level as well.

**Figure 5.**
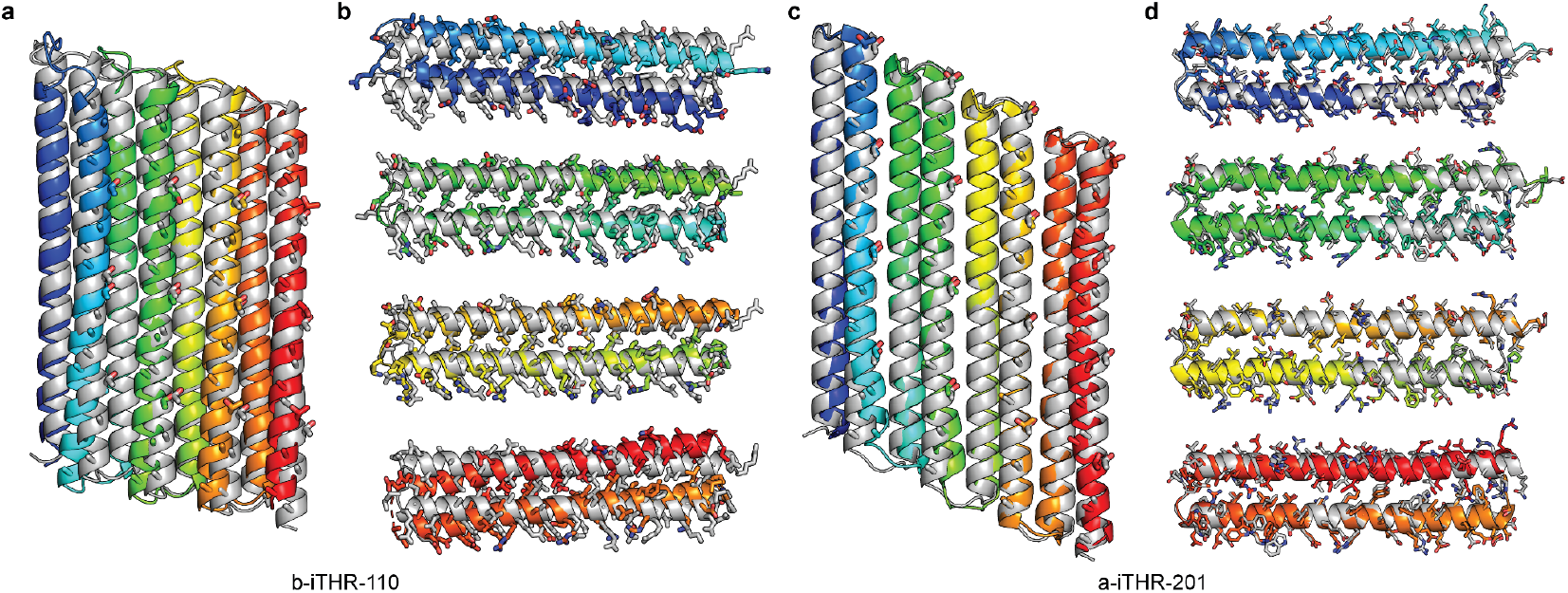
Crystal structures of iTHRs aligned to parametric designed models. **a)** Overlay of the b-iTHR-110 (PDB ID: 9MG8) crystal structure (coloured) and the parametric designed model (grey). Cα-RMSD = 2.5 Å (over 368 atoms). The putative ice-binding threonine and alanine residues are shown as sticks. **b)** Side-views of individual b-iTHR-110 repeats showing all side chains as sticks. **c)** Overlay of the a-iTHR-201 (PDB ID: 9D01) crystal structure (coloured) and the parametric designed model (grey). Cα-RMSD = 1.1 Å (over 370 atoms). The putative ice-binding threonine and alanine residues are shown as sticks. **d)** Side- views of individual a-iTHR-201 repeats showing all side chains as sticks.

## Discussion

We have demonstrated how to design Ice-binding <u>T</u>wistless <u>H</u>elical <u>R</u>epeat (iTHR) proteins, a new topology of ice-binding proteins containing parallel alpha-helices presenting TXXXAXXXAXX motifs, which have IRI activities comparable to those of native IRI-active ice-binding proteins (**Fig. 2h**). The iTHR proteins were designed to target the {110} or {201} facets of ice, and three of the four ice crystal polymorphs are as expected (**Fig. S3**). The remaining design, a-iTHR-110, which appears to stabilize the off-target {201} facet (**Fig. S3a, e**), has the largest spacing between helices and is the least thermostable design (**Fig. 2c**), perhaps indicating that the off-target binding is due to poorer core packing and greater flexibility. Variants of the designs were used to show that geometric complementarity to ice-crystal surfaces alone is sufficient to obtain significant IRI activity (**Fig. 3**). We observed that increasing the size of the ice-binding surface does not translate into a higher IRI activity (**Fig. 4**). The IRI activity of flexible antifreeze glycoproteins and polyvinyl alcohol polymers increases with increasing chain length^28^, however, for native ice-binding beta-roll repeat proteins it has been established that the relation between ice-binding and number of repeats is non-monotonic, with a clear optimal length^34,35^. Some authors have attributed this to a slight mismatch in spacing, helix bending, and/or misfolding.

The IRI activities of these iTHRs are more than 1 order of magnitude higher than the previously designed TIP ice-binders^20^. Many (but not all) natural IRI-active proteins also have thermal hysteresis (TH) activity^1,2^; in fact, some of the most potent IRI-active proteins display little or no TH activity. We have found that none of the iTHR designs have measurable TH activity, not even at concentrations of 3 mM (unpublished). Currently, the design of proteins with TH activity is challenging, since it is not yet clear what physical properties are required for proteins to show TH activity. Super-resolution microscopy results suggest that TH activity requires long residence times of ice-binding proteins on the ice surface^36^, whereas proteins with short residence times only show IRI activity. Performing similar super-resolution microscopy experiments on the designed TIP and iTHR proteins may inform the design of TH-active proteins. Additionally, as the iTHR proteins are highly stable–especially compared to many native ice-binding proteins–we can systematically vary the interfacial ‘X’ residues in the TXXXAXXXAXX motifs and the geometry and chemistry of the non-IBS surface and measure how these changes affect IRI and TH activity.

The present iTHR designs may offer advantages for applications of ice-binding proteins that are currently being explored such as cryopreservation of cells, tissues, and organs^37^, since they are thermostable and express with high yield. The highly regular and modular structures of these iTHRs may allow further engineering of the ice-water interface by incorporating additional protein-protein interfaces^16^ to form large protein assemblies that drive ice nucleation, or protein nanocages that encapsulate ice crystals. Additionally, recent advances in deep-learning methods, such as RFdiffusion^38^, may enable the design of ice-binding sites and motifs that are not seen in nature.

## Materials & Methods

### Computational design

An ice-binding helix (“B” helix) with N = 2 or 4 copies of the putative ice-binding motif TXXXAXXXAXX with a minor helical pitch of ω_1_ = 98.2° per residue was generated using a python script that incorporates Watson-Crick parameters^18,39^. The “B” helix was aligned to the z-axis by its helical axis and centred at the origin in PyMOL. Next, the “B” helix was rotated about the z-axis such that the Cα atom to Cβ atom vector of the threonine is pointing parallel to the y-axis. Subsequently, the phase of the “B” helix is sampled in 18 increments of 6° to create diversity in the orientation of the ice-binding residues facing the solvent.

Next, a stabilizing helix (“S” helix) — with ω_1_ = 100° per residue — was generated using the BundleGridSampler^40^ with 96 varieties of geometry defined by a subtag of the mover: <Helix r0=“10” delta_omega0_min=“52” delta_omega0_max=“68” delta_omega0_samples=“3” z0_offset_min=“6” z0_offset_max=“10” z0_offset_samples=“4” invert=“true” delta_omega1_min=“1” delta_omega1_max=“96” delta_omega1_samples=“8”/>. For each “B” helix and “S” helix pair, rigid bodies are copied and translated using the RigidBodyTransMover in Pyrosetta following the translations in **Table S1**. Residues that were not “T” or “A” motif residues were mutated to “G” to simplify tracking of residue identities downstream.

The rigid body translated copies of the “B” and “S” helix pairs were then looped together with the ConnectChainsMover in RosettaScripts. Because this looped unit now contains (i) two copies of each helix, (ii) all the necessary loops built for a repeat protein, and (iii) the intended transformation between repeat units, a repeat protein model can be built from this unit that can be extended arbitrarily with the RepeatPropagationMover^16^; in this case we chose to move forward with 4 repeat units so that the interior 2 repeats could be identical in sequence, representing the potentially duplicatable interface between repeats..

At this point, all residue identities in the models are either “T” or “A” if they are part of the ice-binding motif, or “G” if they are a non-motif part of the helix, or “V” if they were built with the ConnectChainsMover. ProteinMPNN^25^ was used to design all of the residues on these structures except for positions that were already “T” or “A”; and each of the 4 repeats was required to design the same sequence as the other repeats. “C” was the only prohibited residue identity. All residues between each repeat (asymmetric unit) were linked, except for solvent-accessible residues at the capping repeats to avoid forming hydrophobic caps.

Subsequently,structures were predicted from the designed sequences using both AlphaFold2^26^ and RoseTTAFold^27^. Designs with < 2.5 Å Cα root mean square deviation (Cα-RMSD) between the AlphaFold2 model and the parametrically generated backbone and < 5 Å RMSD between the AlphaFold2 and RoseTTAFold predictions were selected for experimental characterization.

### Synthetic Gene Bacterial Protein Expression and Purification

Synthetic gene fragments encoding 36 iTHR designs were codon optimized for expression in Escherichia coli were acquired from IDT and cloned into a vector pET-29(b+)-derived vector, using golden gate assembly. The constructs were transformed into BL21 (DE3) *E. coli* competent cells.

Transformants were cultured in 4×1 mL Terrific Broth medium supplemented with 200 mgmL^-1^ kanamycin in deep 96-well plates. Expression, under the control of a T7 promoter, proceeded for 24 h at 37 °C using Studier autoinduction^41^. Cells pellets were resuspended in lysis buffer (50 mM Tris pH 8.0, 300 mM NaCl, 30 mM Imidazole, with 1 mg of DNAseI) and chemically lysed by addition of BugBuster® (Sigma-Aldrich). The soluble fraction, clarified by centrifugation, underwent purification via Ni^2+^ immobilised metal affinity chromatography (IMAC) using Ni-NTA resin. The resin with bound cell lysate was washed with ten column volumes of wash buffer (50 mm Tris, pH 8.0 + 300 mM NaCl, 30 mM imidazole) followed by elution with 50 mM Tris pH 8.0 + 300 mM NaCl, 300 mM imidazole. The soluble fractions were subjected to SDS–polyacrylamide gel electrophoresis analysis.

Expression cultures were scaled up to 0.5 L for further characterization, with expression again proceeding for 24 h at 37 °C using 1 mM IPTG at OD 0.6 before harvesting by centrifugation and purification in similar conditions. After purification proteins were loaded on a size exclusion chromatography Superdex Increase 200 10/300 gl (Cytiva) column in running buffer PBS pH 7.4. Purified protein was checked for correct mass via Liquid Chromatography–Mass Spectrometry (**Table S3**)

### Liquid Chromatography–Mass Spectrometry Measurements (LC-MS)

Molecular weights of each protein were determined by reverse-phase liquid chromatography-mass spectrometry (LC-MS) with an Agilent G6230B time-of-flight (TOF) instrument on an AdvanceBio RP-Desalting column. The solvents used were water with 0.1% formic acid (Solvent A) and acetonitrile with 0.1% formic acid (Solvent B). The spectra were then deconvoluted with a total entropy algorithm in BioConfirm.

### MW-assisted solid-phase peptide synthesis (MW-SPPS)

The *wf*AFP was synthesized on TentaGel™ S-RAM resin (loading 0.25 mmol/g, 100 µm), following the Fmoc/t-Bu strategy under nitrogen atmosphere. The sequence elongation was performed on a 0.1 mmol scale with a five-fold molar excess of Fmoc-AA-OH on a microwave-assisted solid-phase peptide synthesizer (Liberty Blue, CEM, Matthews, NC, U.S.A.). Activation was achieved with DIC and Oxyma Pure at 90°C. The reaction temperatures were monitored by an internal fiber-optic sensor and deprotection and coupling reactions were performed in a Teflon vessel, applying microwave energy under nitrogen bubbling. The peptide elongation was performed by repeating the MW-SPPS cycle for each amino acid, using a modified protocol based on the CARBOMAX method provided by CEM. wfAFP sequence: *H-***D**TASDAAAAAA**L**TAANAKA AAE**L**TAANAAAAAA**A**TAR*-NH*_*2*_

In short, the peptide was elongated with the following cycles. Fmoc deprotection was performed using a 20% (v/v) piperidine solution in DMF in 4 steps: 25°C, 2W, 30s, *ΔT*=5°C; 75°C, 125W, 20s, ΔT=1°C; 90°C, 5W, 5s, ΔT=0°C; 90°C, 20W, 90s, ΔT=0°C, followed by washing with DMF (3x 4mL). Coupling was performed with Fmoc-AA-OH (2.5 mL, 0.2 M in DMF, 0.5 mmol, 5 eq), DIC (1 mL, 1.0 M in DMF, 1 mmol, 10 eq) and Oxyma Pure (0.5 mL, 1.0 M in DMF, 0.5 mmol, 5 eq) in 4 steps: 40°C, 5W, 30s, ΔT=5°C; 75°C, 163W, 12s, ΔT=1°C; 90°C, 5W, 9s, ΔT=0°C; 90°C, 20W, 120s, ΔT=0°C, followed by a DMF wash step (4mL). All couplings after β-branched amino acids (in bold) used prolonged coupling times of 240 seconds for the final step.

Once the synthesis was complete, the resin was transferred into a syringe equipped with a filter, washed with DMF (3×4mL), DCM (3×4mL) and dried under nitrogen flow. Crude peptide was obtained after acidic deprotection and cleavage from resin with TFA/TIS/H_2_O (10 mL; 190/5/5; v/v/v) for two hours while shaking, followed by precipitation in cold Et_2_O (50 mL). Purification by RP-HPLC (linear-gradient 10-70% ACN in water with 0.1% TFA, 60 min, ReproSil Gold 120 C18, 10 µm - 240 × 20 mm, 5mL/min) yielded *wf*AFP as a white powder after lyophilization in 8.59 mg (2.65 µmol). HPLC-MS analysis was performed to validate purity, using a C18 Jupiter SuC4300A 150 × 2.0 mm column using miliQ water with 0.1% FA and acetonitrile with 0.1% FA, using a gradient of 5% to 100% ACN over 10 minutes, connected to a Thermo Fisher LTQ XL Linear Ion Trap Mass Spectrometer (rt = 3.25 min) (**Fig. S4**). Exact mass calculated [C_133_H_226_N_44_O_50_ + H]^+^ : 3241.6595; [C_133_H_226_N_44_O_50_ + 2H]^2+^ : 1621.3334; [C_133_H_226_N_44_O_50_ + 3H]^3+^ : 1081.2247; [C_133_H_226_N_44_O_50_ + 4H]^4+^ : 811.1703.

### Circular Dichroism Measurements

Proteins were diluted to 0.2 mg/mL in PBS (10 mM sodium phosphate pH 7.4, 140 mM NaCl) in a quartz cuvette with a 1 mm pathlength. On a JASCO J-1500 (JASCO Corporation), Spectral scans from 200 to 280 nm were performed at 25°C, 65°C and 95°C where temperature was ramped at rate of 1 °C/min and ellipticity was monitored at 222 nm.

### Ice Recrystallization Inhibition Assay

Samples were prepared by dilution of the protein to the indicated concentration in PBS + 40 wt% sucrose. Subsequently, a 1 µL sample was applied on a 22 × 22-mm coverslip, and a second coverslip was lowered on top of the drop so that the sample was sandwiched between the coverslips and transferred to a Nikon ECLIPSE Ci-Pol optical microscope equipped with a Nikon L Plan 20× (NA 0.45) objective and the Linkam LTS420 stage. This stage was controlled by the Linksys32 software. To measure the ice-recrystallization rates, the sample was first completely frozen by reducing the temperature to −40 °C with 20 °C/min. After freezing, the temperature was gradually increased to −10 °C with 10 °C/min and then further to −8 °C with 1 °C/min, upon which individual crystals could be observed and the sample was stabilised. Recrystallization was monitored by obtaining an image each minute. IRI rates were then analysed using ImageJ and Python. In brief, the 8-bit images of the ice crystals were subjected to the bandpass filter, enhance contrast and subtract background functions of imageJ. Subsequently, the bright signal at the edges and then the individual crystals were isolated by the autoThreshold function using the Otsu method, followed by the Convert to Mask function. “Analyze particles” was then used to obtain the area of each crystal. These data were imported into python, upon which the radius was calculated to determine the cubed number-average radius per image. Recrystallization growth rates were determined by applying a linear fit to the resulting cubed number-average radius as a function of time traces

### Ice Shaping Assay

To monitor ice shaping in the presence of the various proteins, the samples were prepared similarly to the IRI assays, but diluted into PBS containing 20 wt% sucrose. The sample was stabilized around −4.3 °C, so that a few crystals were observed in the field of view. After stabilization, the samples were supercooled with decreasing temperature 0.2 °C/min, which forced the crystals to shape in presence of the proteins, or PBS buffer only. Samples were monitored with 1 or 5 s intervals using a Nikon 50× ELWD objective. Stills that were used in the figures were taken after 1 min of supercooling (full views in **Fig. S3**).

### Crystallization

Crystallization samples were prepared by concentrating a-iTHR-201 protein to 52 mg/mL and b-iTHR-110 to 14.3 mg/mL in TBS (20 mM Tris-HCl pH 8, 150 mM NaCl). All crystallization experiments were conducted using the sitting drop vapor diffusion method. Crystallization trials were set up in 200 nL drops using the 96-well plate format at 20 °C. Crystallization plates were set up using a Mosquito from SPT Labtech, then imaged using UVEX microscopes and UVEX PS-600 from JAN Scientific. Diffraction quality crystals formed in 2.8 M Sodium acetate trihydrate pH 7.0 for a-iTHR-201 and 0.2 M Sodium chloride, 28% (w/v) MPD for b-iTHR-110.

Diffraction data was collected at the Advanced Light Source (ALS) HHMI beamline 8.2.1/ 8.2.2. X-ray intensities and data reduction were evaluated and integrated using XDS^42^ and merged/scaled using Pointless/Aimless in the CCP4 program suite^42,43^. Starting phases were obtained by molecular replacement using Phaser^44^ using the designed model for the structures. Following molecular replacement, the models were improved using phenix.autobuild^45^. Structures were refined in Phenix^45^. Model building was performed using COOT^46^. The final model was evaluated using MolProbity^47^. Data collection and refinement statistics are recorded in **Table S4**. Data deposition, atomic coordinates, and structure factors reported in this paper have been deposited in the Protein Data Bank (PDB)^48^, http://www.rcsb.org/, with accession code: 9D01 (a-iTHR-201) and 9MG8 (b-iTHR-110).

## Acknowledgements

This work was financially supported by the Dutch Research Council to I.K.V (NWO-VICI: 232.106), the European Research Council to I.K.V (ERC-2020-CoG 101001965), the VLAG Graduate School Research Fellowship and Fulbright Visiting Scholar Fellowship to R.J.d.H. H.P. was supported by the Open Philanthropy Project Improving Protein Design Fund. T.F.H. was supported by the DARPA MIMA Seedling grant (HR00112420369) and by the Breakthrough Fund 100 nanometer Assemblies program from The Audacious Project. A.K.B., A.C., A.K., and E.J. were supported by The Bill and Melinda Gates Foundation, #INV-010680. E.B. was supported by the GR014929: C19 NIH PAN-COV P01-62-7718. D.B. was supported by The Howard Hughes Medical Institute. We thank the Advanced Light Source (ALS) beamline 8.2.1/8.2.2 at Lawrence Berkeley National Laboratory for X-ray crystallographic data collection. The Berkeley Center for Structural Biology is supported by the NIH, National Institute of General Medical Sciences, and the HHMI. The ALS is supported by the Director, Office of Science, Office of Basic Energy Sciences and US Department of Energy (DOE) (DE-AC02-05CH11231). We thank Tim Hogervorst for providing the wfAFP peptide.

## Data availability

All data and design models are available in the main text, the Supplementary Information or https://doi.org/10.5281/zenodo.13763849. Crystallographic datasets with resolved side chains have been deposited in the Protein Data Bank (PDB ID: 9MG8 for b-iTHR-110; PDB ID: 9D01 for a-iTHR-201).

## Code availability

An example RosettaScripts script and input for generating iTHR proteins are provided at https://doi.org/10.5281/zenodo.13763849. The example script was confirmed to successfully run with PyRosetta version 4 as available at https://pyrosetta.org (ref. 49). Documentation for ProteinMPNN sequence design is available at https://github.com/dauparas/ProteinMPNN (ref. 25). Designs were filtered with AlphaFold 2 available at https://github.com/google-deepmind/alphafold (ref. 26).

## Notes

### Competing Interest Statement

The authors have declared no competing interest.

## References

1. Bar Dolev, M., Braslavsky, I. & Davies, P. L. Ice-Binding Proteins and Their Function. Annu Rev Biochem 85, 515–542 (2016).

2. Olijve, L. L. C. et al. Blocking rapid ice crystal growth through nonbasal plane adsorption of antifreeze proteins. Proc. Natl. Acad. Sci. U. S. A. 113, 3740–3745 (2016).

3. Budke, C., Heggemann, C., Koch, M., Sewald, N. & Koop, T. Ice recrystallization kinetics in the presence of synthetic antifreeze glycoprotein analogues using the framework of LSW theory. J. Phys. Chem. B 113, 2865–2873 (2009).

4. Kristiansen, E. & Zachariassen, K. E. The mechanism by which fish antifreeze proteins cause thermal hysteresis. Cryobiology 51, 262–280 (2005).

5. Lin, M., Cao, H. & Li, J. Control strategies of ice nucleation, growth, and recrystallization for cryopreservation. Acta Biomater. 155, 35–56 (2023).

6. Kristiansen, E. et al. Hyperactive antifreeze proteins from longhorn beetles: some structural insights. J. Insect Physiol. 58, 1502–1510 (2012).

7. Bouvet, V. & Ben, R. N. Antifreeze glycoproteins. Cell Biochemistry and Biophysics 39, 133–144 (2003).

8. Harding, M. M., Anderberg, P. I. & Haymet, A. D. J. ‘Antifreeze’ glycoproteins from polar fish. European Journal of Biochemistry 270, 1381–1392 (2003).

9. Huang, P.-S., Boyken, S. E. & Baker, D. The coming of age of de novo protein design. Nature 537, 320–327 (2016).

10. Ueda, G. et al. Tailored design of protein nanoparticle scaffolds for multivalent presentation of viral glycoprotein antigens. Elife 9, (2020).

11. Hsia, Y. et al. Design of multi-scale protein complexes by hierarchical building block fusion. Nat Commun 12, 2294 (2021).

12. Doyle, L. et al. Rational design of α-helical tandem repeat proteins with closed architectures. Nature 528, 585–588 (2015).

13. Pyles, H., Zhang, S., De Yoreo, J. J. & Baker, D. Controlling protein assembly on inorganic crystals through designed protein interfaces. Nature 571, 251–256 (2019).

14. Davila-Hernandez, F. A. et al. Directing polymorph specific calcium carbonate formation with de novo protein templates. Nat. Commun. 14, 8191 (2023).

15. Saragovi, A. et al. Controlling semiconductor growth with structured de novo protein interfaces. bioRxiv 2024.06.24.600095 (2024) doi:10.1101/2024.06.24.600095.

16. Huddy, T. F. et al. Blueprinting extendable nanomaterials with standardized protein blocks. Nature 627, 898–904 (2024).

17. Sicheri, F. & Yang, D. S. C. Ice-binding structure and mechanism of an antifreeze protein from winter flounder. Nature 375, 427–431 (1995).

18. de Haas, R. J. et al. De novo designed ice-binding proteins from twist-constrained helices. Proceedings of the National Academy of Sciences 120, e2220380120 (2023).

19. Kwan, A. H.-Y. et al. Solution structure of a recombinant type I sculpin antifreeze protein. Biochemistry 44, 1980–1988 (2005).

20. de Haas, R. J. et al. De novo designed ice-binding proteins from twist-constrained helices. Proceedings of the National Academy of Sciences 120, e2220380120 (2023).

21. And, P. D. & Sönnichsen*, F. D. Source of the Ice-Binding Specificity of Antifreeze Protein Type I. (2000) doi:10.1021/ci000449b.

22. Bowles, D. J., Lillford, P. J., Rees, D. A. & Shanks, I. A. Structure and function of antifreeze proteins. Philosophical Transactions of the Royal Society of London. Series B: Biological Sciences (2002) doi:10.1098/rstb.2002.1081.

23. Goto, A., Hondoh, T. & Mae, S. The electron density distribution in ice Ih determined by single-crystal x-ray diffractometry. J. Chem. Phys. 93, 1412–1417 (1990).

24. Huang, P.-S. et al. RosettaRemodel: a generalized framework for flexible backbone protein design. PLoS One 6, e24109 (2011).

25. Dauparas, J. et al. Robust deep learning-based protein sequence design using ProteinMPNN. Science 378, 49–56 (2022).

26. Jumper, J. et al. Highly accurate protein structure prediction with AlphaFold. Nature 596, 583–589 (2021).

27. Baek, M. et al. Accurate prediction of protein structures and interactions using a three-track neural network. Science 373, 871–876 (2021).

28. Budke, C. et al. Quantitative Efficacy Classification of Ice Recrystallization Inhibition Agents. (2014) doi:10.1021/cg5003308.

29. Bar, M., Celik, Y., Fass, D. & Braslavsky, I. Interactions of β-Helical Antifreeze Protein Mutants with Ice. (2008) doi:10.1021/cg800066g.

30. The effects of steric mutations on the structure of type III antifreeze protein and its interaction with ice. Journal of Molecular Biology 275, 515–525 (1998).

31. de Haas, R. J. et al. Flat Solenoidal Ice-Binding Proteins as Scaffolds for Solid-Binders. Advanced Materials Interfaces 10, 2300001 (2023).

32. Antifreeze Protein from Freeze-Tolerant Grass Has a Beta-Roll Fold with an Irregularly Structured Ice-Binding Site. Journal of Molecular Biology 416, 713–724 (2012).

33. Zhang, D.-Q., Liu, B., Feng, D.-R.He, Y.-M. & Wang, J.-F. Expression, purification, and antifreeze activity of carrot antifreeze protein and its mutants. Protein Expr Purif 35, 257–263 (2004).

34. Marshall, C. B., Daley, M. E., Sykes, B. D. & Davies, P. L. Enhancing the Activity of a β-Helical Antifreeze Protein by the Engineered Addition of Coils†. Biochemistry (2004) doi:10.1021/bi0488909.

35. Naullage, P. M., Qiu, Y. & Molinero, V. What Controls the Limit of Supercooling and Superheating of Pinned Ice Surfaces? The Journal of Physical Chemistry Letters (2018) doi:10.1021/acs.jpclett.8b00300.

36. Tas, R. P., Hendrix, M.M.R.M. & Voets, I. K. Nanoscopy of single antifreeze proteins reveals that reversible ice binding is sufficient for ice recrystallization inhibition but not thermal hysteresis. Proceedings of the National Academy of Sciences 120, e2212456120 (2023).

37. [No title]. EMBO Press 10.15252/embr.202052162.

38. Watson, J. L. et al. De novo design of protein structure and function with RFdiffusion. Nature 620, 1089–1100 (2023).

39. Huang, P.-S. et al. High thermodynamic stability of parametrically designed helical bundles. Science 346, 481–485 (2014).

40. Dang, B. et al. De novo design of covalently constrained mesosize protein scaffolds with unique tertiary structures. Proc. Natl. Acad. Sci. U. S. A. 114, 10852– 10857 (2017).

41. Studier, F. W. Protein production by auto-induction in high density shaking cultures. Protein Expr. Purif. 41, 207–234 (2005).

42. Kabsch, W. XDS. Acta Crystallogr. D Biol. Crystallogr. 66, 125–132 (2010).

43. Winn, M. D. et al. Overview of the CCP4 suite and current developments. Acta Crystallogr. D Biol. Crystallogr. 67, 235–242 (2011).

44. McCoy, A. J. et al. Phaser crystallographic software. J. Appl. Crystallogr. 40, 658–674 (2007).

45. Adams, P. D. et al. PHENIX: a comprehensive Python-based system for macromolecular structure solution. Acta Crystallogr. D Biol. Crystallogr. 66, 213–221 (2010).

46. Emsley, P. & Cowtan, K. Coot: model-building tools for molecular graphics. Acta Crystallogr. D Biol. Crystallogr. 60, 2126–2132 (2004).

47. Williams, C. J. et al. MolProbity: More and better reference data for improved all-atom structure validation. Protein Sci. 27, 293–315 (2018).

48. Berman, H. M. et al. The Protein Data Bank. Nucleic Acids Res. 28, 235–242 (2000).

49. Chaudhury, S., Lyskov, S. & Gray, J. J. PyRosetta: a script-based interface for implementing molecular modeling algorithms using Rosetta. Bioinformatics 26, 689–691 (2010).

