## Supplementary material for "Inhibition of ice recrystallization with designed twistless helical repeat proteins": SI

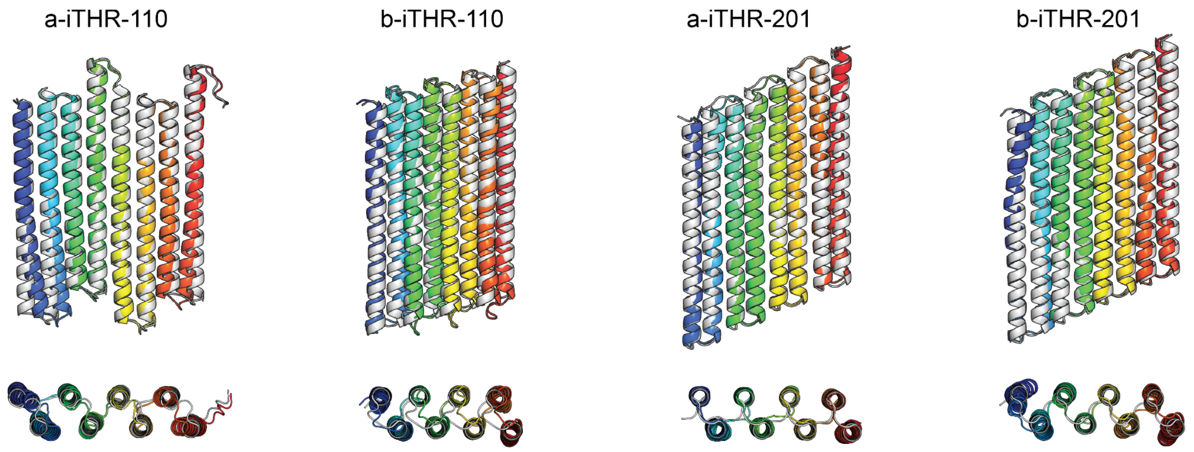

**Figure S1.** Alignments of AlphaFold2 vs. parametrically designed models for a-iTHR-110, b-iTHR-110, a-iTHR-201, and b-iTHR-201 (RMSD = 1.27 Å, 1.62 Å, 1.07 Å, 1.44 Å, respectively).

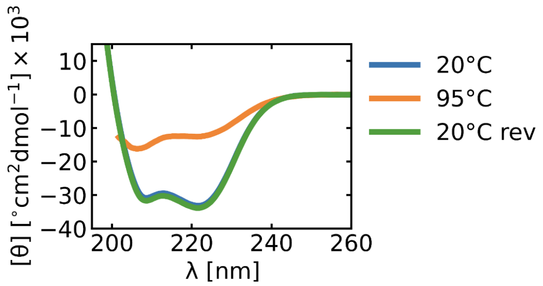

**Figure S2.** Circular dichroism of a-iTHR-110 with spectra at 20°C, 95°C and back to 20°C (20°C rev). At 95°C the signal is reduced significantly but when cooled back to 20°C a-iTHR-110 has identical signal, suggesting refolding.

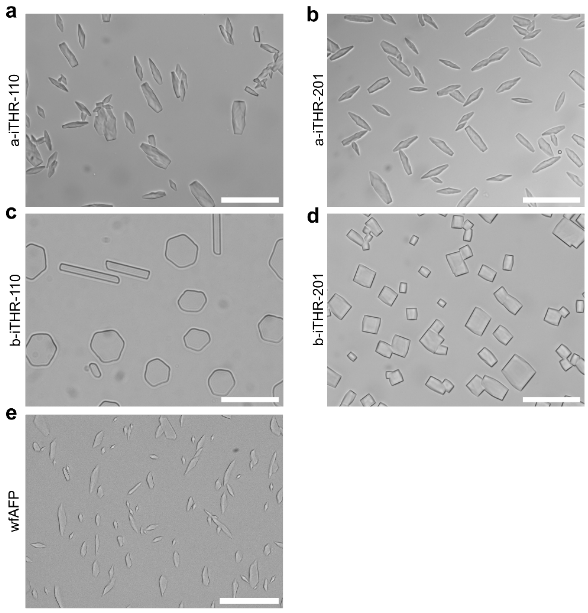

**Figure S3. Full views of ice shaping experiments**.

**a)** a-iTHR-110 at 113.3 uM **b)** a-iTHR-201 at 58.8 uM **c)** b-iTHR-110 at 26.0 uM **d)** b-iTHR-201 at 101.7 uM **e)** wfAFP at 20.0 uM. Stills shown after 60 seconds of cooling at 0.2 °C per minute. Range of cooling was set to -4.3°C to -4.5°C, except a-iTHR-201 (-4.5°C to -4.7°C). The expected ice crystal shapes for {110} binders were hexagonal and rectangular, reflecting the facets, while bipyramidal or blunt-end bipyramidal shapes were anticipated for {201} binders. This trend holds true except for a-iTHR-110, which shows bipyramidal and blunt-end shaping. Shaping for the {201}-binders is consistent with shaping observed for wfAFP, reported to bind to the {201} facet. Scale bar: 100 µm.

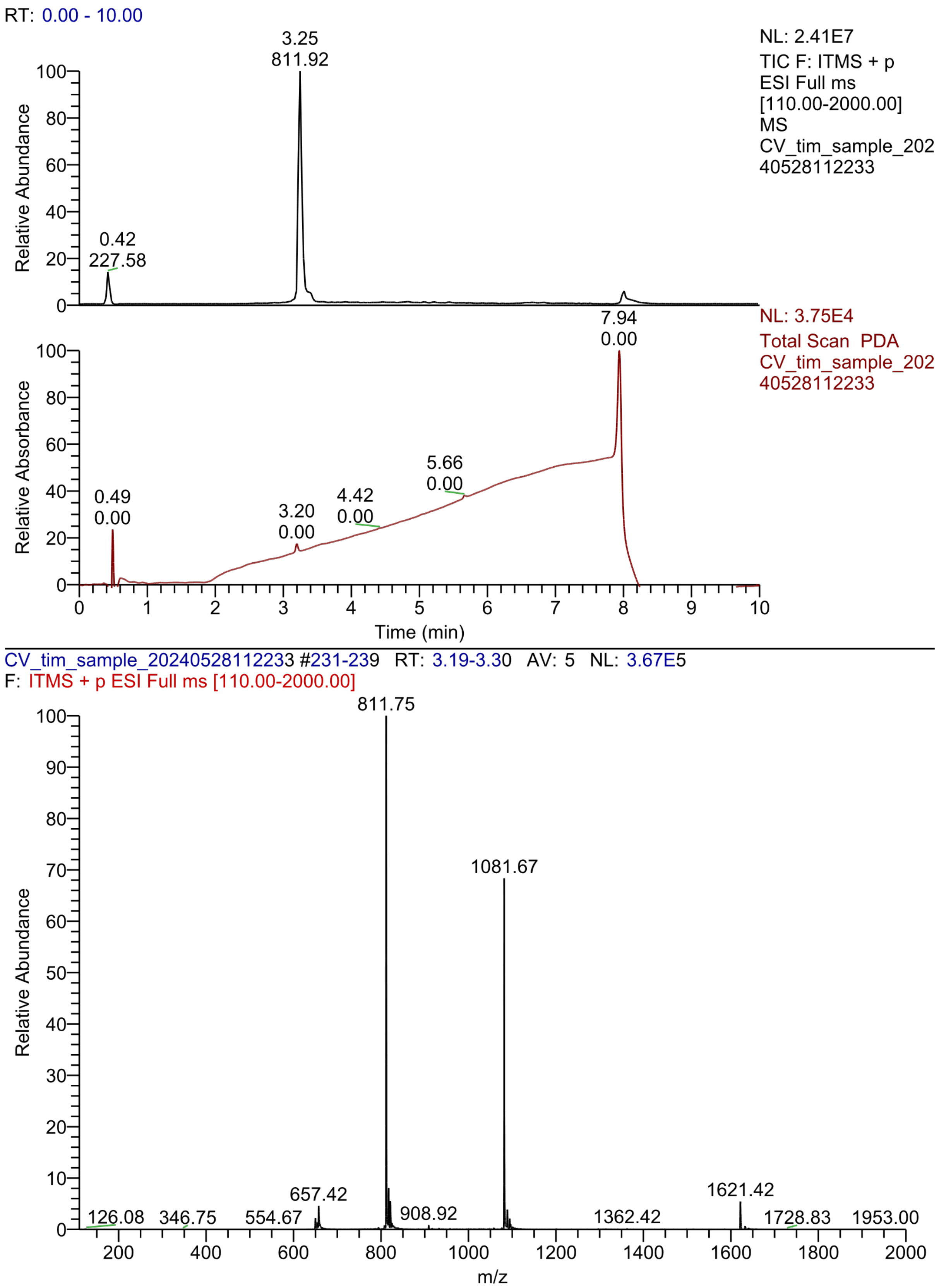

**Figure S4.** HPLC-MS wfAFP (rt = 3.25 min) 5-100% ACN in MQ (+0.1% formic acid).

**Table S1.** Translations identified for matching oxygen spacings on {110} plane hexagonal ice using twist constrained helices with threonine spacing of 16.7Å. The internal symmetric repeat can be 1 or 2 ice-binding helices. Translations shown are relative to the first ice-binding helix ($B_0$).

| ID | helices per repeat | ΔX ($B_0$ → $B_1$) | ΔZ ($B_0$ → $B_1$) | ΔX ($B_0$ → $B_2$) | ΔZ ($B_0$ → $B_2$) | Target facet |
|---|---|---|---|---|---|---|
| 1 | 1 | -10.375 | 2.828 | -20.751 | 5.656 | {110} |
| 2 | 1 | 10.375 | -2.828 | 20.751 | -5.656 | {110} |
| 3 | 2 | -10.375 | 2.828 | -20.75 | -10.954 | {110} |
| 4 | 2 | 10.375 | -2.828 | 20.75 | 10.954 | {110} |
| 5 | 2 | -10.376 | 19.438 | -20.752 | 22.266 | {110} |
| 6 | 2 | 10.376 | -19.438 | 20.752 | -22.266 | {110} |
| 7 | 1 | -13.832 | -7.354 | -27.663 | -14.708 | {110} |
| 8 | 1 | 13.832 | 7.353 | 27.663 | 14.707 | {110} |
| 9 | 2 | -13.833 | 9.256 | -27.664 | 1.902 | {110} |
| 10 | 2 | 13.833 | -9.257 | 27.664 | -1.903 | {110} |
| 11 | 1 | 10.375 | 2.828 | 20.751 | 5.656 | {110} |
| 12 | 1 | -10.375 | -2.828 | -20.751 | -5.656 | {110} |
| 13 | 2 | 10.375 | 2.828 | 20.752 | -10.954 | {110} |
| 14 | 2 | -10.375 | -2.828 | -20.752 | 10.954 | {110} |
| 15 | 2 | 10.374 | 19.438 | 20.75 | 22.266 | {110} |
| 16 | 2 | -10.374 | -19.438 | -20.75 | -22.266 | {110} |
| 17 | 1 | 13.832 | -7.354 | 27.663 | -14.708 | {110} |
| 18 | 1 | -13.832 | 7.353 | -27.663 | 14.707 | {110} |
| 19 | 2 | 13.831 | 9.256 | 27.662 | 1.902 | {110} |
| 20 | 2 | -13.831 | -9.257 | -27.662 | -1.903 | {110} |

**Table S1 Continued.** Translations identified for matching oxygen spacings {201} planes hexagonal ice using twist constrained helices with threonine spacing of 16.7Å. The internal symmetric repeat can be 1 or 2 ice-binding helices. Translations shown are relative to the first ice-binding helix ($B_0$).

| ID | Helices per repeat | ΔX ($B_0 \rightarrow B_1$) | ΔZ ($B_0 \rightarrow B_1$) | ΔX ($B_0 \rightarrow B_2$) | ΔZ ($B_0 \rightarrow B_2$) | Target facet |
|---|---|---|---|---|---|---|
| 21 | 1 | 8.264 | 3.678 | 16.529 | 7.356 | {201} |
| 22 | 1 | -8.264 | -3.679 | -16.529 | -7.357 | {201} |
| 23 | 2 | 8.264 | 3.678 | 16.53 | -9.254 | {201} |
| 24 | 2 | -8.264 | -3.679 | -16.53 | 9.253 | {201} |
| 25 | 1 | -8.265 | 13.01 | -16.53 | 26.019 | {201} |
| 26 | 1 | 8.264 | -13.01 | 16.529 | -26.019 | {201} |
| 27 | 1 | -8.264 | -3.601 | -16.528 | -7.202 | {201} |
| 28 | 1 | 8.263 | 3.6 | 16.527 | 7.201 | {201} |
| 29 | 2 | -8.264 | -3.6 | -16.529 | 9.409 | {201} |
| 30 | 2 | 8.263 | 3.6 | 16.528 | -9.409 | {201} |
| 31 | 1 | -8.264 | 3.678 | -16.529 | 7.356 | {201} |
| 32 | 1 | 8.264 | -3.679 | 16.529 | -7.357 | {201} |
| 33 | 2 | -8.264 | 3.678 | -16.528 | -9.254 | {201} |
| 34 | 2 | 8.264 | -3.679 | 16.528 | 9.253 | {201} |
| 35 | 1 | 8.265 | 13.01 | 16.53 | 26.019 | {201} |
| 36 | 1 | -8.264 | -13.01 | -16.529 | -26.019 | {201} |
| 37 | 1 | 8.266 | -3.601 | 16.532 | -7.202 | {201} |
| 38 | 1 | -8.265 | 3.6 | -16.531 | 7.201 | {201} |
| 39 | 2 | 8.266 | -3.6 | 16.531 | 9.409 | {201} |
| 40 | 2 | -8.265 | 3.6 | -16.53 | -9.409 | {201} |
| 57 | 1 | 12.396 | 5.516 | 24.792 | 11.032 | {201} |
| 58 | 1 | 12.396 | -5.516 | 24.792 | -11.032 | {201} |
| 59 | 1 | -12.396 | 5.516 | -24.792 | 11.032 | {201} |
| 60 | 1 | -12.396 | -5.516 | -24.792 | -11.032 | {201} |
| 61 | 1 | 12.396 | 5.516 | 24.792 | -5.668 | {201} |
| 62 | 1 | 12.396 | -5.516 | 24.792 | 5.668 | {201} |
| 63 | 1 | -12.396 | 5.516 | -24.792 | -5.668 | {201} |
| 64 | 1 | -12.396 | -5.516 | -24.792 | 5.668 | {201} |

**Table S2.** LC-MS theoretical mass vs. experimental mass

| Name | Theoretical mass (kDa) | Experimentally determined mass (kDa) |
|---|---|---|
| **a-iTHR-110** | 44.619 | 44.619 |
| **b-iTHR-110** | 40.624 | 40.624 |
| **a-iTHR-201** | 44.321 | 44.321 |
| **b-iTHR-201** | 43.247 | 43.247 |

**Table S3.** Sequences of proteins generated in this study.

| Name | Sequence |
|---|---|
| **a-iTHR-110** | MSGRLAEETEEQIKELEEKEKERQEELEELLREAEEEAQKQLEKEAGDPAAR<br>KAILTRLRAEQEAVLTERRASQEATYTERLALLEATYTRGALEPERYIELLREL<br>EEWYLEQLKELLERAREQARRLLELGRGDPAVEEAVLTWLEAEQKAVETQK<br>RAVLQATETRIRAILEAWETRRGVRLGERLAEETERQIEELRKLEEERQEQLE<br>ELLRLAEEEAKRQLELEAGDPAARKAILTRLRAEQEAVLTERRASQEATYTER<br>LALLEATYTRGALEPERYIELLEELEEWYQEQLDELLKRAEEQAKRLKELGRG<br>DPAVEKAVETWLEAEEKAVKTQKEAVAEATKTRIQAILEAWETRRGVRLGEG<br>SGSHHWGSTHHHHHH |
| **b-iTHR-110** | MSGVPPEELLKGAEEFIEELIRESEEGAEALLQALEEAIEAAEEAARRKSGTGE<br>EVGAALAAAVTEVIAALSALLTETLAHVAALATQALAAAAAQRVPPEELLKGAE<br>RFIELLIRLSERGAEALLRALELAIEAAEEAARRKSGTGKEVGAALAAAVTEVIA<br>ALSALLTLTLAHVAALATQALAAAAAQRVPPEELLRGAERFIELLIRLSERGAE<br>ALLRALELAIRAAREAARRKSGTGEEVGAALAAAVTEVIAALSALLTLTLAHVA<br>ALATQALAAAAAQRVPPEELEKGAERFIELLERLSERGAEALERALELAEEAA<br>EEAARRKSGTGEEVGAALAAAETEVEAALSALETLTEAHVAALATQAEAAAA<br>AQRGSGSHHWGSTHHHHHH |
| **a-iTHR-201** | MSGAPSPESARRQLAELEETQERGEAEIARAVEEGERELERLEEEQKAGLPA<br>TEFHARAIAALTEWIARQEALLTRLQAELEAQFTRLLAELKAGLEPSRETLEQF<br>LELREFWQEQGERQLALARELTDRQLEWVKRLKEWGLPQTEANALAIATLTR<br>GAALLEALLTRTVAGVEAIFTEALAALRAGLEPSRETLEQFLELREFWQEQGE<br>RQLALARELTDRQLEWVKRLKEWGLPQTEANALAIATLTRGAALLEALLTRTV<br>AGVEAIFTEALAALRAGLEPSEEFTELLEELRERFVEAYERLKEIARELTEWLL<br>RWVEEFREWGLPETEAEALREAVETRGRALLEAIETRGEAAVEAIRTEREAA<br>RRAGQGSGSHHWGSTHHHHHH |
| **b-iTHR-201** | MSGLPPREWFERAKEEIERVREREEEELEELKREWEEAREEAKERLERGEP<br>VTEVTAWLRAAATEVIARREAVVTRALAERRALITEALARVRAAGLPWEELLA<br>AAREAIELLREAGEWALEAIREILELAREAAKELLELGADQTEVSAFLRAQATR<br>AAALQEAVVTETFAQQVAIITEALAAVTAAGLPWEELLAAAREAIELLREAGEW<br>ALEAIREILELAREAAKELLELGADQTEVSAFLRAQATRAAALQEAVVTETFAQ<br>QVAIITEALAAVTAAGVPPEEFRRAAEEAIEWVEETEKWALEQIEEILRLAEEA<br>AAFLREIGTDETEVEAFLEAARTRAEALAEAVKTETEALKEAIRTEAEAYVAAA<br>EGSGSHHWGSTHHHHHH |
| **a-iTHR-110.KO$^{Ala}$** | MSGRLAEETEEQIKELEEKEKERQEELEELLREAEEEAQKQLEKEAGDPAAR<br>KAILTRLREEQEAVLTERRESQEATYTERLELLEATYTRGALEPERYIELLREL<br>EEWYLEQLKELLERAREQARRLLELGRGDPAVEEAVLTWLEEEQKAVETQK<br>REVLQATETRIREILEAWETRRGVRLGERLAEETERQIEELRKLEEERQEQLE<br>ELLRLAEEEAKRQLELEAGDPAARKAILTRLREEQEAVLTERRESQEATYTER<br>LELLEATYTRGALEPERYIELLEELEEWYQEQLDELLKRAEEQAKRLKELGRG<br>DPAVEKAVETWLEEEEKAVKTQKEEVAEATKTRIQEILEAWETRRGVRLGEG<br>SGSHHWGSTHHHHHH |
| **a-iTHR-110.KO$^{Thr}$** | MSGRLAEETEEQIKELEEKEKERQEELEELLREAEEEAQKQLEKEAGDPAAR<br>KAILERLRAEQEAVLEERRASQEATYEERLALLEATYERGALEPERYIELLREL<br>EEWYLEQLKELLERAREQARRLLELGRGDPAVEEAVLEWLEAEQKAVEEQK<br>RAVLQATEERIRAILEAWEERRGVRLGERLAEETERQIEELRKLEEERQEQLE<br>ELLRLAEEEAKRQLELEAGDPAARKAILERLRAEQEAVLEERRASQEATYEER<br>LALLEATYERGALEPERYIELLEELEEWYQEQLDELLKRAEEQAKRLKELGRG<br>DPAVEKAVEEWLEAEEKAVKEQKEAVAEATKERIQAILEAWEERRGVRLGEG<br>SGSHHWGSTHHHHHH |
| **a-iTHR-110x0.5** | MSGRLAEETEEQIKELEEKEKERQEELEELLREAEEEAQKQLEKEAGDPAAR<br>KAILTRLRAEQEAVLTERRASQEATYTERLALLEATYTRGALEPERYIELLEEL |

| | |
|---|---|
| | EEWYQEQLDELLKRAEEQAKRLKELGRGDPAVEKAVETWLEAEEKAVKTQK<br>EAVAEATKTRIQAILEAWETRRGVRLGEGSGSHHWGSTHHHHHH |
| **a-iTHR-110x2** | MSGRLAEETEEQIKELEEKEKERQEELEELLREAEEEAQKQLEKEAGDPAAR<br>KAILTRLRAEQEAVLTERRASQEATYTERLALLEATYTRGALEPERYIELLREL<br>EEWYLEQLKELLERAREQARRLLELGRGDPAVEEAVLTWLEAEQKAVETQK<br>RAVLQATETRIRAILEAWETRRGVRLGERLAEETERQIEELRKLEEERQEQLE<br>ELLRLAEEEAKRQLELEAGDPAARKAILTRLRAEQEAVLTERRASQEATYTER<br>LALLEATYTRGALEPERYIELLRELEEWYLEQLKELLERAREQARRLLELGRG<br>DPAVEEAVLTWLEAEQKAVETQKRAVLQATETRIRAILEAWETRRGVRLGER<br>LAEETERQIEELRKLEEERQEQLEELLRLAEEEAKRQLELEAGDPAARKAILTR<br>LRAEQEAVLTERRASQEATYTERLALLEATYTRGALEPERYIELLRELEEWYL<br>EQLKELLERAREQARRLLELGRGDPAVEEAVLTWLEAEQKAVETQKRAVLQA<br>TETRIRAILEAWETRRGVRLGERLAEETERQIEELRKLEEERQEQLEELLRLAE<br>EEAKRQLELEAGDPAARKAILTRLRAEQEAVLTERRASQEATYTERLALLEAT<br>YTRGALEPERYIELLEELEEWYQEQLDELLKRAEEQAKRLKELGRGDPAVEK<br>AVETWLEAEEKAVKTQKEAVAEATKTRIQAILEAWETRRGVRLGEGSGSHH<br>WGSTHHHHHH |
| **b-iTHR-201x0.5** | MSGLPPREWFERAKEEIERVREREEEELEELKREWEEAREEAKERLERGEP<br>VTEVTAWLRAAATEVIARREAVVTRALAERRALITEALARVRAAGVPPEEFRR<br>AAEEAIEWVEETEKWALEQIEEILRLAEEAAAFLREIGTDETEVEAFLEAARTR<br>AEALAEAVKTETEALKEAIRTEAEAYVAAAEGSGSHHWGSTHHHHHH |
| **b-iTHR-201x2** | MSGLPPREWFERAKEEIERVREREEEELEELKREWEEAREEAKERLERGEP<br>VTEVTAWLRAAATEVIARREAVVTRALAERRALITEALARVRAAGLPWEELLA<br>AAREAIELLREAGEWALEAIREILELAREAAKELLELGADQTEVSAFLRAQATR<br>AAALQEAVVTETFAQQVAIITEALAAVTAAGLPWEELLAAAREAIELLREAGEW<br>ALEAIREILELAREAAKELLELGADQTEVSAFLRAQATRAAALQEAVVTETFAQ<br>QVAIITEALAAVTAAGLPWEELLAAAREAIELLREAGEWALEAIREILELAREAA<br>KELLELGADQTEVSAFLRAQATRAAALQEAVVTETFAQQVAIITEALAAVTAAG<br>LPWEELLAAAREAIELLREAGEWALEAIREILELAREAAKELLELGADQTEVSA<br>FLRAQATRAAALQEAVVTETFAQQVAIITEALAAVTAAGLPWEELLAAAREAIE<br>LLREAGEWALEAIREILELAREAAKELLELGADQTEVSAFLRAQATRAAALQE<br>AVVTETFAQQVAIITEALAAVTAAGLPWEELLAAAREAIELLREAGEWALEAIR<br>EILELAREAAKELLELGADQTEVSAFLRAQATRAAALQEAVVTETFAQQVAIIT<br>EALAAVTAAGVPPEEFRRAAEEAIEWVEETEKWALEQIEEILRLAEEAAAFLRE<br>IGTDETEVEAFLEAARTRAEALAEAVKTETEALKEAIRTEAEAYVAAAEGSGS<br>HHWGSTHHHHHH |
| **wfAFP** | DTASDAAAAAALTAANAKAAAELTAANAAAAAAATAR |

**Table S4.** X-ray crystallography data collection and refinement statistics.

| | a-iTHR-201 (9D01) | b-iTHR-110 (9MG8) |
|---|---|---|
| **Data collection** | | |
| Space group | P 43 21 2 | P21 |
| Cell dimensions | | |
| *a, b, c* (Å) | 64.24, 64.24, 188.32 | 42.91, 75.59, 115.99 |
| α, β, **γ** (°) | 90, 90, 90 | 90.00, 90.31, 90.00 |
| Resolution (Å) | 47.08 - 2.18 (2.25 - 2.18) | 46.01 - 2.50 (2.60 - 2.50) |
| $R_{merge}$ | 0.07 (1.76) | 0.124 (1.571) |
| **I / σI** | 20.3 (19.9) | 7.1 (1.1) |
| Completeness (%) | 99.9 (99.0) | 97.9 (97.7) |
| Redundancy | 19.5 (19.9) | 5.0 (5.4) |
| **Refinement** | | |
| Resolution (Å) | 44.90 - 2.18 (2.25 - 2.18) | 46.01 - 2.5 (2.58 - 2.5) |
| No. reflections | 21415 (1357) | 24486 (2097) |
| $R_{work}$ / $R_{free}$ | 0.2587 (0.4555)/ 0.3122 (0.5245) | 0.2644 (0.3182)/ 0.3092 (0.3650) |
| No. atoms | | |
| Protein | 2972 | 5124 |
| Water | 13 | 13 |
| B-factors (Å$^2$) | | |
| Protein | 82 | 83 |
| Water | 66 | 67 |
| r.m.s. deviations | | |
| Bond lengths (Å) | 0.003 | 0.004 |
| Bond angles (°) | 0.461 | 0.584 |
